# *In-silico* modeling and interpretation of RBP binding disentangle m^6^A-RBP interaction

**DOI:** 10.1101/2024.11.23.624962

**Authors:** Jianche Liu, Xinlu Zhu, Yang Yin, Zhoutong Xu, Jialin He, Xushen Xiong

## Abstract

RNA binding protein (RBP) binding and *N*^6^-methyladenosine (m^6^A) are both essential post-transcriptional regulatory layers for RNA fate decisions. However, the intricate mechanism underlying the interaction between m^6^A and RBP binding remains underexplored. Here, we develop TransRBP, an interpretable deep learning framework, to model the base-resolution binding of RBPs from RNA sequences and to subsequently investigate the interaction between m^6^A and RBPs. TransRBP achieves a median accuracy of 0.59 across 32 m^6^A-related RBPs, representing a 28% increase over the state-of-the-art model. Using gradient-based interpretation, we demonstrate that the binding motifs of the m^6^A-related RBPs strongly enrich for splicing consensus, laying a foundation for studying the RBP-dependent crosstalk between m^6^A and splicing. Moreover, we develop an *in-silico* mutagenesis assay to assess the impact of m^6^A on RBPs, and utilize the self-attention mechanism to elucidate the interplay between RBP binding and m^6^A. We further uncover 1,806 variant-RBP combinations with the *in-silico* mutagenesis, revealing variants that strongly alter RBP binding for genetic diseases including Parkinson’s disease, autism, and cardiomyopathy. In particular, we identify m^6^A *cis-*acting variants that alter RBP binding in an m^6^A-proximal manner, including the binding of UPF1 that contributes to Alzheimer’s disease, and the DDX3X binding to cardiomyopathy and muscular dystrophy. Together, TransRBP accurately models the binding of RBP and its interaction with m^6^A, shedding light on the m^6^A-RBP dynamics and providing multi-layer mechanistic insights for genetic diseases.

## Introduction

The regulatory code of genomic sequence extends beyond the process of transcription. Post-transcriptional regulation mechanisms at various RNA levels have profound impacts on gene expression^1,2,3^. One of the key mechanisms is the regulation of RNA-binding proteins (RBPs), which orchestrate nearly every aspect of RNA fate decision and metabolism^3,4^. The binding specificity of an RBP is jointly determined by short sequence motif and RNA structure, which can be affected by both local and distal sequences^5^. Moreover, modifications in RNAs, such as methylation and pseudouridylation, can alter these interactions and lead to changes in RNA stability, localization, and splicing patterns, providing an additional layer of regulation^6,7^.

Among over 170 types of RNA modifications discovered, N^6^-methyladenosine (m^6^A) has been established as the most prevalent and functionally important mRNA modification in humans^5,8^. The interplay between RBPs and m^6^A plays a crucial role in the dynamic regulation of the RNA life cycle. RBP could act as m^6^A writers (METTL3, METTL14, etc.) and adaptors (RBM15 and WTAP) to deposit m^6^A modifications, or erasers (FTO and ALKBH5) to remove m^6^A modifications, while m^6^A can influence RBP readers (e.g. the YTH family) to regulate various aspects in RNA metabolisms^5,8^.

The interplay between m^6^A and relevant RBPs is essential in a wide range of physiological and pathological processes, including embryonic and tissue development, cancer progression, cell differentiation, neuron functions and various human diseases^9–14^.

Despite the well-established interactions between m^6^A and its corresponding RBP regulators, emerging studies have gradually indicated more complex and interconnected interactions between m^6^A and a broader range of RBPs^15–19^. The findings have largely expanded the regulatory landscape of m^6^A and deepened the disease mechanisms dependent on such m^6^A-RBP interactions. These RBPs may be directly affected by m^6^A or act as co-factors with the primary m^6^A regulators to interact with m^6^A. For example, FMR1 was recently recognized as a novel m^6^A reader RBP, whose phase switch is instructed by m^6^A and contributes to mRNA decay^18^. SRSF3 and SRSF10, previously identified as splicing factors, were also reported to be recruited or modulated for splicing in a m^6^A-YTHDC1-dependent manner^19^. However, regardless of the large variety of eukaryotic RBPs and the continuing realization of their interplay with m^6^A, a comprehensive repertoire of m^6^A-interacted RBPs and a systematic understanding of the interaction dynamics remains largely incomplete.

Genome-wide association studies (GWASs) have laid important foundations for the association between genetic variants and complex human diseases^20–24^. However, over 95% of the GWAS variants do not alter protein sequences, complicating the mechanisms of their regulatory roles in disease pathogenesis^25–27^. Genetic studies have been extensively focused on interpreting variant impact on the mRNA expression levels^28–30^. However, the post-transcriptional regulations have been largely neglected, leading to critical mechanistic gaps in the functional output of genes given the discrepancy between transcription and protein expression^31,32^. Specifically, the effects of these disease variants on RBP binding, and the potential interactions between variants, m^6^A, and RBPs remain poorly understood. Investigating the molecular basis of these disease variants at the post- transcriptional regulatory layers would lead to a more comprehensive understanding of genetic diseases.

An initial step toward answering these questions is to precisely measure the RBP binding profile. Large-scale experimental techniques such as cross-linking and immunoprecipitation followed by sequencing (CLIP-seq) and its derivatives^33^, including enhanced CLIP (eCLIP^34^), have provided high-resolution datasets for RBP binding profiling. However, the identified sequence determinants of RBP binding specificity are very limited so far, likely due to the technical challenges and the high costs of the CLIP-seq experiments^34,35^. Moreover, given the inherent limitations of experimental data in extrapolating RBP binding on unseen RNA sequences, comprehensive identification of essential regulatory codes intrinsic to RBP binding remains challenging. Therefore, computational modeling may serve as an important complementary tool for imputing the binding of RBPs and extracting missing binding information embedded in the RNA sequences.

Drawing parallels with natural language processing (NLP), deep learning holds great potential to leverage genomic sequences as inputs for modeling molecular phenotypes, interpreting the ’languages’ of genes^36–39^. However, existing deep learning models for predicting RBP binding from the sequence were primarily framed as classification tasks, that merely determine whether a region is bound or unbound, without the resolution of predicting the actual binding profile of RBPs^40,41,42^.

While RBPnet^43^ has been developed to model the binding profile, it only processed a short receptive field of sequence (300 bp) using convolutional modules. This may largely limit its ability in utilizing the distal information, thereby losing the opportunity to capture the influence of tertiary RNA structure or the interactions with RNA modifications during RBP modeling. Moreover, efforts to develop the interpretation toolsets for elucidating how RNA modifications, such as m^6^A, interact with RBPs, are currently lacking.

In this study, we introduce TransRBP, a Transformer-based deep learning framework for modeling RBP binding profiles at a base resolution from RNA sequences. With a substantially increased input sequence length and the inclusion of attention modules, our model outperforms the state-of-the-art sequence-to-signal RBP model. Based on TransRBP, we further develop integrated tools for model interpretation. Utilizing the gradient-based interpretation algorithms, we reveal that m^6^A-related RBP binding motifs are notably enriched in splicing consensus, providing a foundation to interrogate the RBP-dependent crosstalk between m^6^A and splicing. Further, we develop an *in-silico* mutagenesis assay to define an impact score of m^6^A on RBPs, and utilize the self-attention mechanism to elucidate the interplay between RBPs and m^6^A. We assess the impact of more than 18,000 genetic disease variants on the binding of m^6^A-related RBPs, spanning 24 complex diseases relevant to brain, heart, lung, and muscle tissues. We further reveal 1,806 variant-RBP combinations, where the disease variants strongly alter RBP binding. Importantly, we identify m^6^A *cis*-acting disease variants that impact RBP binding in a distance-to-m^6^A manner, shedding light on the molecular mechanisms for genetic diseases, including the binding of UPF1 on Alzheimer’s disease, and the binding of DDX3X on cardiomyopathy and muscular dystrophy.

## Results

### Prioritization of 32 m^6^A-related RBPs

To better model the interaction between m^6^A and RBPs, we first prioritized RBPs with binding site enrichments of m^6^A loci. We gathered the genome-wide binding profiles of 139 RBPs from ENCODE^44,45^, which were based on eCLIP experiments (**Methods**, **Supplementary Table S1**). To identify RBPs with a preferred binding of m^6^A-modified RNA, we performed an enrichment analysis of the intersection between RBP binding sites and m^6^A modifications profiled in the same cell line^46,47^. On average, each of the 139 RBPs has about 37 binding sites overlapped with m^6^A peaks, representing approximately 1% of their total peaks (**Supplementary Table S2**).

Based on the quantification of intersection, we prioritized 32 m^6^A-related RBPs whose binding sites are among the top 50 m^6^A-intersected number and the top 50 m^6^A-intersected ratio (**Fig. 1a**). These 32 RBPs on average exhibited about 129 binding sites intersecting with m^6^A peaks, which correspond to about 2.8% of their total peak numbers (**Extended Data Fig. 1a**). Intriguingly, these RBPs formed a tight protein-protein interaction (PPI) network with the established m^6^A readers, writers, adapters, and erasers (**Fig. 1b**). The tight interconnection network suggests that these RBPs may be involved in m^6^A regulation^16^, and therefore largely expanding the known repertoire of m^6^A regulators. Consistently, 18 of these RBPs (56%) possess a well-defined RNA binding domain (**Extended Data Fig. 1b**), and are predominantly involved in the regulation of RNA stability, mRNA metabolism, and translation (**Extended Data Fig. 1c**).

**Figure 1.**
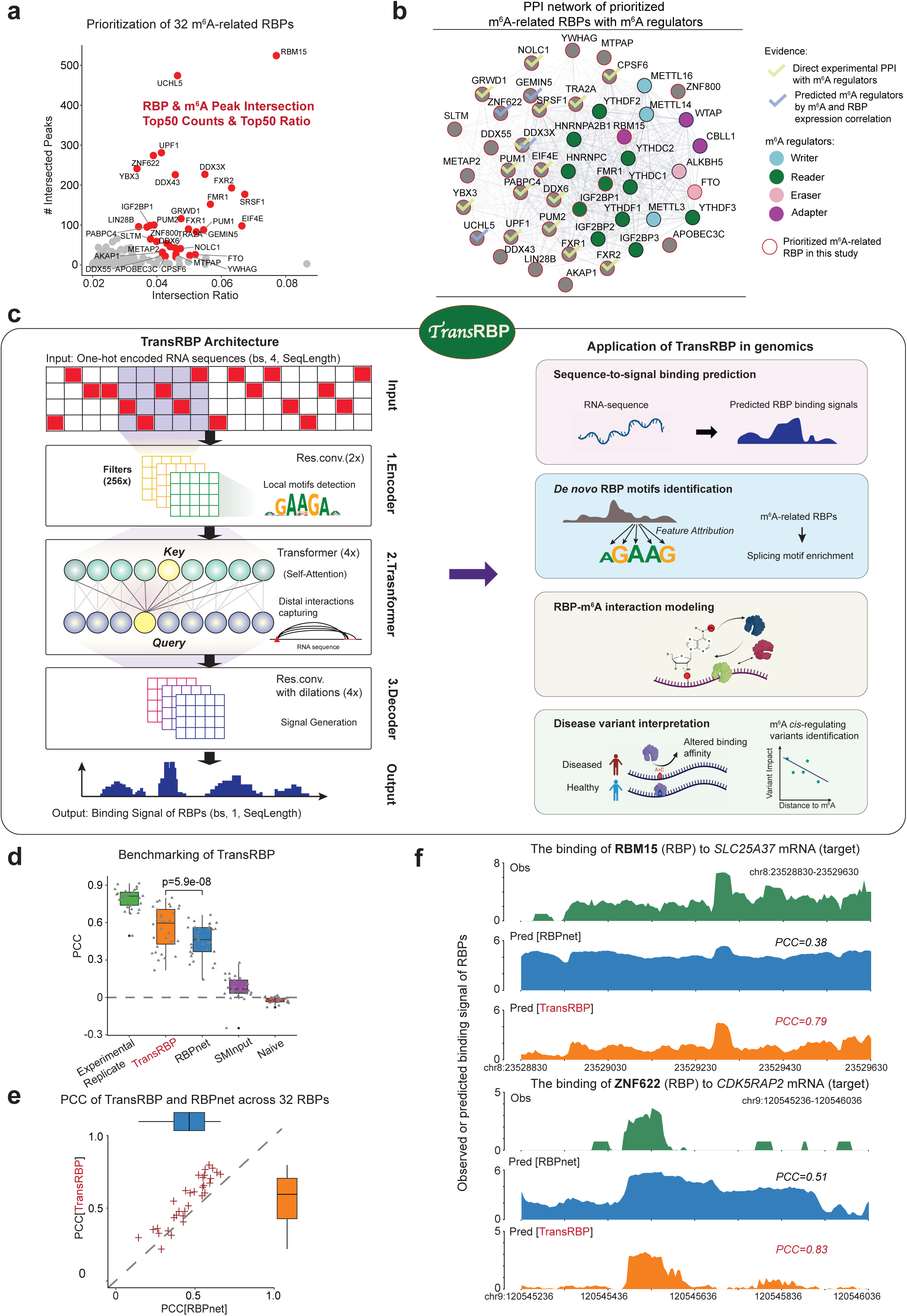
TransRBP accurately models the binding of m^6^A-related RBPs. **(a)** Scatter plot showing the number of m^6^A-intersected peaks (y-axis) and the m^6^A-intersected ratio (x-axis) of the RBPs collected from ENCODE. RBPs having both the top 50 m^6^A-intersection number and ratio are prioritized as m^6^A-related RBPs and are colored in red. **(b)** Protein-protein interaction (PPI) network of the prioritized m^6^A-related RBPs (red circle) and the known m^6^A regulators (in different colors based on their types). The prioritized RBPs supported by experimentally confirmed direct interactions with m^6^A regulators and by RBP binding sites enrichment in m^6^A QTLs^16^ are highlighted in yellow and blue check marks, respectively. **(c)** The architecture of the convolution-attention model (TransRBP) trained for predicting RBP binding signals from 800 bp RNA sequences (left panel), and the multiscaled interpretation of TransRBP (right panel). The application and interpretation include binding prediction from sequences, *de novo* RBP motif discovery from feature attribution, RBP-m^6^A interaction modeling, and disease variant interpretation by *in-silico* mutagenesis. **(d)** Pearson correlation coefficient (PCC) between the observed and predicted RBP binding profile evaluated on the held-out test set for TransRBP and RBPnet^43^, and compared to experimental replicates, size-matched input (SMInput) and randomly initialized untrained model (Naive). TransRBP is highlighted in red. **(e)** Comparison of the PCCs between the RBPnet (x-axis) and TransRBP (y-axis) across the 32 m^6^A-related RBPs. **(f)** Examples of observed (Obs) and predicted (Pred) RBP binding profiles of SRSF1 on *SLC25A37* mRNA (top), and ZNF622 binding on *CDK5RAP2* mRNA (bottom) in the test set. PCCs between the observed and predicted tracks were shown at the top-right corner of each panel.

### TransRBP accurately models RBP binding from sequence

To understand the underlying binding syntax of the m^6^A-related RBPs, we designed TransRBP, an interpretable deep learning framework for the base-resolution RBP binding prediction using RNA sequence as the input.

We employed a convolution-attention-coupled architecture in TransRBP to learn biologically relevant motifs embedded in sequence and meanwhile capture potential interactions between these motifs^48,49^ (**Fig. 1c** and **Extended Data Fig. 1d**, **Methods**). The model starts with a pre-convolution layer composed of 256 convolutional filters, with each filter spanning 5 bp of sequence, to extract local motifs derived from the sequence^48,50^. An exponential activation function is subsequently employed to facilitate a robust motif derivation from the gradient-based feature attribution^51^ (**Extended Data Fig. 1d**). The model encoder then uses two layers of residual CNN to promote information propagation. After encoding, the sequence embeddings are processed through a 4-layer Transformer block, which enhances the model’s ability in learning complex interactions between sequences via the self-attention mechanism^48^. Finally, a dilated CNN is used as the decoder for RBP binding profile generation, which helps in increasing the receptive field and effectively maintains the spatial organization of the input bases. The model architecture of TransRBP was designed and fine-tuned based on an extensive evaluation of various network architectures and hyperparameters (**Extended Data Fig. 2a**-c). Notably, we utilized a receptive field of 800 bp, which is substantially larger than the width of 100 bp or 300 bp used by existing RBP binding prediction models^40,42,43^. This optimization significantly increases the accuracy of RBP prediction by incorporating more sequence information (**Extended Data Fig. 2c**), and meanwhile enables the model to capture the potential long-distance interaction within the RNA sequence.

We developed a sliding-window-based data augmentation method to expand the quantity and diversity of the dataset, and to ensure that the binding signal is evenly distributed in the 800 bp sequence (**Extended Data Fig. 2d**, **Methods**). This can effectively prevent the model from overfitting to the shape of RBP binding profile rather than learning the sequence-dependent information. For each RBP, a total of 20,000 sequences were randomly selected from the augmented dataset, and were then split by chromosome into training, validation, and test sets (**Methods**). To further account for the experimental biases of eCLIP caused by UV radiation or contamination^33^, we paired each eCLIP experimental data to its size-matched input (SMInput) control data to normalize the binding profile of the RBPs and to mitigate the impact of unspecific binding (**Methods**).

We next systematically evaluated the performance of our model. Across the 32 m^6^A-related RBPs, our model achieved a median Pearson correlation coefficient (PCC) of 0.59 and a median Spearman correlation coefficient (SCC) of 0.53 between the predicted and the observed RBP binding profiles (**Fig. 1d**, **Extended Data Fig. 2e**). We then carried out benchmarking by comparing our model to RBPnet^43^, the state-of-the-art sequence-to-signal model for RBP binding profile prediction. RBPnet is composed of purely deep convolutional neural networks and utilizes multiple residual convolutional layers to model the RBP binding profile from the sequence^43^. Both our model and RBPnet demonstrated higher accuracies in RBP binding prediction compared to the SMInput and Naive models with the randomly initialized parameters (**Fig. 1d, Extended Data Fig. 2e**).

Overall, our model achieved significantly higher accuracies compared to RBPnet (P=5.9 × 10^-8^), based on both the PCC and SCC evaluations (**Fig. 1d, Extended Data Fig. 2e**). In addition to the overall comparison, TransRBP outperformed RBPnet in predicting the binding profiles for 29 out of the 32 (∼91%) analyzed m^6^A-related RBPs (**Fig. 1e and Extended Data Fig. 2e**).

To intuitively demonstrate the performance of TransRBP, we selected two RBPs and their binding profile on the target genes as examples for visualization, including RBM15 binding on *SLC25A37* mRNA, and ZNF622 binding on *CDK5RAP2* mRNA (**Fig. 1f**). In both loci, TransRBP achieved more accurate binding predictions (PCC = 0.79 and 0.83, respectively) on the genome tracks compared to RBPnet (PCC = 0.38 and 0.51, respectively), with cleaner binding signal and reduced background noise (**Fig. 1f**). The improvement of TransRBP’s performance is likely due to the enhanced motif extraction capability with more filters in the first convolutional layer, and the higher-order motif interaction modeling enabled by the Transformer layer^48^. These optimization strategies together enhance TransRBP’s predictive accuracy, meanwhile providing an opportunity for the model to decode the complex regulatory syntax underlying RBP binding profiles.

### Gradient-based model interpretation for RBP binding motif interrogation

The binding specificity of the majority of RBPs is primarily driven by motifs formed by specific sequences^43^. Therefore, we next sought to identify the key sequence elements that contribute to the prediction of the m^6^A-related RBP binding through feature attribution algorithms (**Fig. 2a**). We adjusted the integrated gradient (IG)^52^ with gradient correction^52,53^ to assess the contribution score at a per-nucleotide resolution (**Fig. 2a, Methods**). To evaluate the performance of this strategy, we first visualized the contribution score of SRSF1 binding at the *CROCPP2* and *UBR4* loci, as well as the binding of ZNF622 at the *EXOSC10* and *RINT1* loci (**Fig. 2b**). We observed high contributions of AG-riched sequences in SRSF1 binding, and CAAGAA sequences in ZNF622 binding, respectively, which are consistent with their expected binding preference^54,55^ (**Fig. 2b**).

**Figure 2.**
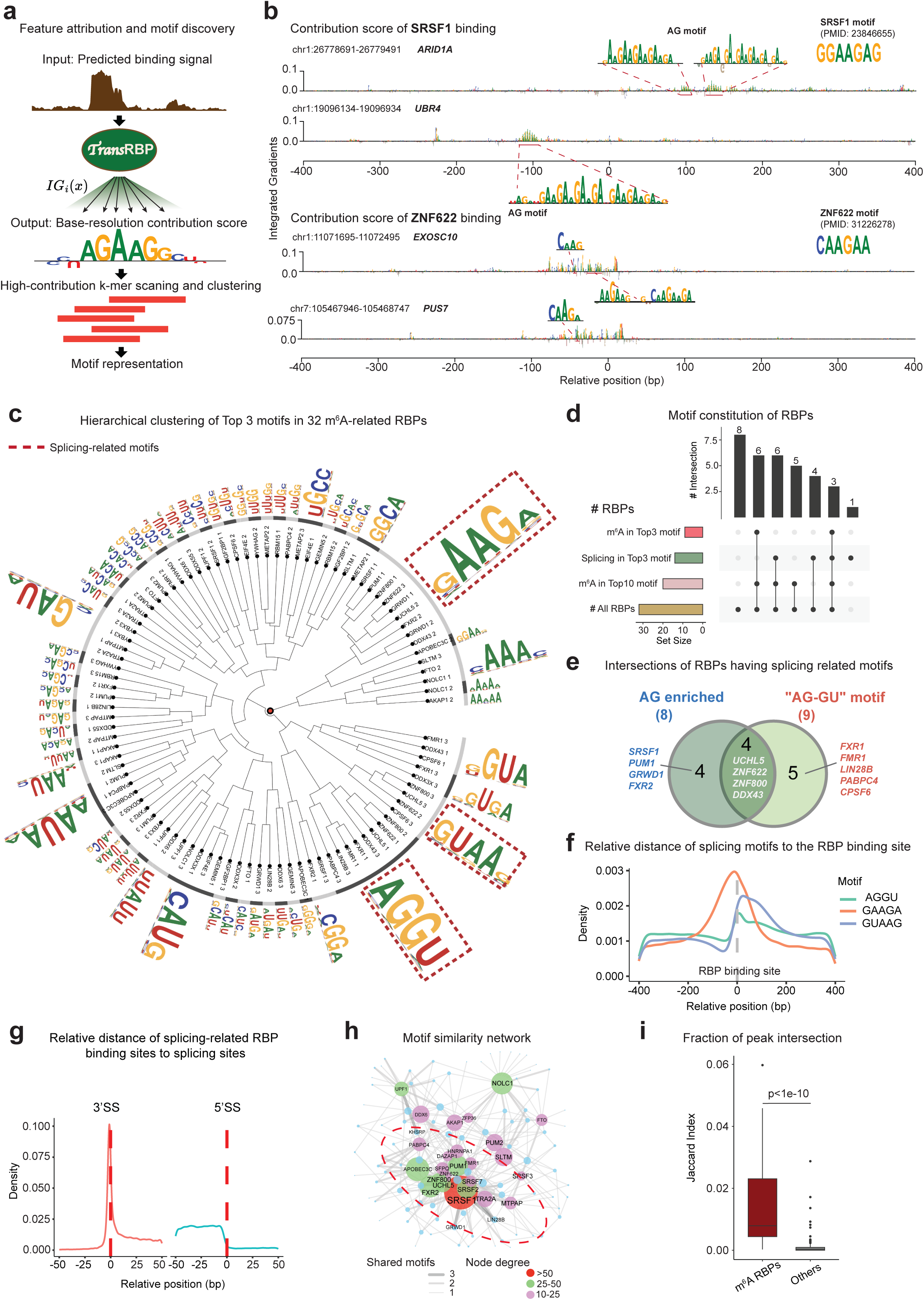
TransRBP enables RBP binding motif discovery using gradient-based interpretation. **(a)** Schema of gradient-based model interpretation from the predicted RBP binding signal for motif discovery by the integrated gradient (IG) algorithm. **(b)** Examples of the contribution scores for SRSF1 binding prediction on *ARID1A* and *UBR4* mRNAs upper), and for ZNF622 binding prediction on *EXOSC10* and *PUS7* (lower). The known motif and the relative reference are shown at the top-right corner of each panel. **(c)** Hierarchical clustering of the PWMs of the top 3 motifs identified from TransRBP interpretation for each m^6^A-related RBP. The three splicing-related motifs were indicated using red dotted boxes. **(d)** Upset plot showing the compositions of 32 m^6^A-related RBPs with regard to their motif types. **(e)** Intersection of RBPs harboring splicing-related AG enriched or AG-GU motifs as their top 3 motifs. **(f)** Averaged distances of *GAAGA*, *AGGU*, and *GUAAG* sequence motifs to the binding sites of the 13 splicing-related RBPs. **(g)** Relative distances of the bindings of the 13 splicing-related RBPs to splicing sites (SS) annotated from the genome on the forward strand. **(h)** Motif similarity network by comparing the top 3 motifs of m^6^A-related RBP with the known motifs of splicing factors reported in the database^56^. Edge weight indicates the number of shared motifs, and the degree of nodes indicates the total number of shared motifs for certain RBPs. **(i)** Jaccard index of peak intersection of SRSF1 with the 32 m^6^A-related RBPs, and with other RBPs on ENCODE. P-value was calculated using the one-way Mann-Whitney U test.

To systematically characterize the binding motifs of m^6^A-related RBPs, we further developed a motif representation strategy that comprehensively scans the base-resolution contribution score along RBP-bound regions using a 5-mer sliding window (**Fig. 2a, Methods**). The resulting high-contribution 5-mers were subsequently clustered into position weight matrices (PWMs) as candidate RBP-binding motifs, and then ranked by the number of supporting sequences for each motif (**Fig. 2a, c, Methods**). Our method identified the RAC (R=A/G) m^6^A consensus as the top motifs in the m^6^A-related RBPs, supporting the involvement of m^6^A in the binding of these RBPs (**Fig. 2d**). In addition, we also confidently recapitulated the known motifs for several well-characterized RBPs, such as the ACUAAC motif for QKI, the GCAUG motif for RBFOX2, and the poly-U motif for HNRNPC^56^ (**Extended Data Fig. 3a**). These results together demonstrated the effectiveness of our interpretation toolkit, providing a reliable method for *de novo* motif identification.

### The binding of m^6^A-related RBPs enrich for splicing consensus

We next sought to understand the global binding preferences of m^6^A-related RBPs. We selected the top 3 motifs for each RBP as the representative motifs (**Extended Data Fig. 3b**), and then performed hierarchical clustering to obtain a comprehensive overview across RBP binding features (**Fig. 2c**). Intriguingly, 13 out of 32 (∼41%) of the RBPs analyzed were preferentially clustered to AG-GU composite motif or AG-enriched motif (**Fig. 2c**). AG and GU are known as the constitutive elements for the 3’ splicing sites (3’SS) and 5’ splicing sites (5’SS)^57,58^, respectively, and the AG- riched motifs are the preferred binding targets of splicing factors^57,58^. We identified 8 RBPs with the AG-riched motif and 9 with the AG-GU motif, respectively, of which 4 RBPs have both motifs identified (**Fig. 2e**). These RBPs included well-established splicing factors and regulators. For example, SRSF1 is a serine/arginine (SR) rich family splicing regulator^57,58^, and LIN28B has been shown to directly regulate the expression of splicing factors and subsequently modulate the global alternative splicing landscape^59^. In addition to the well-established splicing factors, RBPs like FXR1/2, FMR1, and PUM1, which are known for their roles in mRNA transport, stability, and translation^54,55,60^, also show a marked preference for these splicing consensus motifs (**Fig. 2e**), suggesting a potential interplay between splicing and these downstream processes of mRNA. This association is supported by the observed proximity of RBP binding sites to these motifs (**Fig. 2f, Extended Data Fig. 4a**). Moreover, we found RBP binding enrichments of these RBPs surrounding both the splicing junctions (**Fig. 2g, Extended Data Fig. 4b**).

To further explore the potential functional cooperation between m^6^A-related RBPs and known splicing regulators, we designed a network-based motif similarity analysis (**Methods**)^56^. We observed that the primary binding preference of these m^6^A-related RBPs showed extensive sharing of motifs with the motifs of the spliceosome proteins, including SRSF1, SRSF2, and SRSF7^61^, compared to other RBPs (**Fig. 2h, Extended Data Fig. 5a**). Moreover, the binding sites of the SRSF1 show significantly larger fractions of intersection with the binding of m^6^A-related RBPs compared to other RBPs, as indicated by the Jaccard index (**Fig. 2i**). Collectively, the extensively observed association between m^6^A-related RBPs and splicing indicated the potential roles of RBP- dependent m6A regulation in splicing, therefore expanding the regulatory spectrum of the splicing process.

### Model Interpretation disentangles m^6^A-RBP interactions

Since TransRBP is capable of capturing the RBP binding syntax from sequence, we further explored its ability in elucidating the interplay between m^6^A and RBP binding. We developed two complementary model interpretation tools to infer the influence of m^6^A consensus on RBP binding (**Fig. 3a, Methods**). We first quantified the impact of *in-silico* mutations on RBP binding by Kullback- Leibler divergence (KLD) for the sequences at m^6^A-proximal (≤50 bp) and m^6^A-distal regions (>50 bp) in our input, respectively. We then defined the averaged ratio between m^6^A-proximal versus m^6^A-distal KLD to represent the m^6^A-to-RBP impact score, given that the *in-silico* mutations of m^6^A- proximal sequence can be regarded as the interference on m^6^A while m^6^A-distal region can serve as a background estimation (**Fig. 3b, Methods**). The results show that the binding of RBPs including PUM2, PABPC4, FTO, and UCHL5 may undergo strong alteration by m^6^A interference (**Fig. 3b**), with PUM2 showing the most pronounced KLD in the m^6^A-proximal site (**Fig. 3c, Extended Data Fig. 6a**).

**Figure 3.**
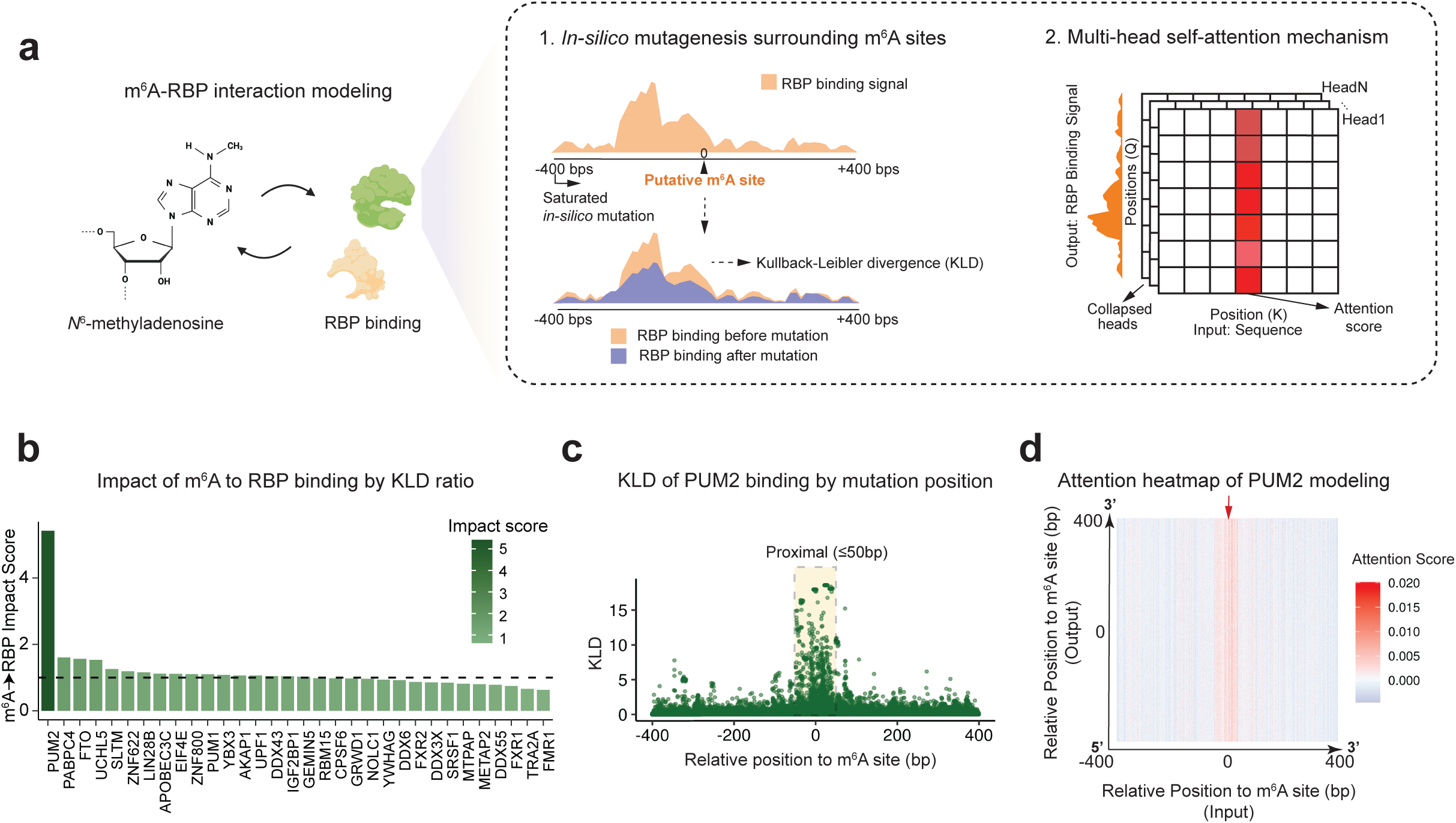
TransRBP disentangles m^6^A-RBP interactions by *in-silico* mutagenesis and self-attention mechanism. **(a)** Schema of *in-silico* mutagenesis and self-attention mechanism for m^6^A-to-RBP impact interpretation based on the TransRBP model. K and Q represent key and query in the Transformer architecture, respectively. **(b)** Quantification of m^6^A-to-RBP impact score across 32 m^6^A-related RBPs by the ratio of average mutation impact (represented by KLD) of m^6^A-proximal (≤50 bp) sites and m^6^A-distal (>50 bp) sites. **(c)** Distribution of mutation impact (KLD) on PUM2 binding with *in-silico* mutagenesis surrounding m^6^A sites (800 bp). The m^6^A-proximal region is denoted by the shaded box. **(d)** Transformer-derived normalized attention weight heatmap surrounding m^6^A sites (800 bp) for PUM2 binding modeling. The arrow indicates the m^6^A site in the input sequence.

As a second metric, we calculated the pairwise positional attention scores from the Transformer architecture to capture the key input positions contributing to model prediction (**Fig. 3a, Methods**). Specifically, the attention weight at position (i, j) reflects the extent to which the i-th element of the output (RBP profile) is influenced by the j-th element of the input (RNA sequence)^48^. We found that the regions surrounding the m^6^A sites showed globally higher attention scores for PUM2 binding prediction (**Fig. 3d, Extended Data Fig. 6b**), in line with the KLD-dependent observation that PUM2 showed the strongest interaction with m^6^A. These results collectively demonstrate the potential of TransRBP, in combination with the *in-silico* interpretation tools, for disentangling the molecular interactions between m^6^A modification and RBP binding.

### RBP-dependent genetic variants of human complex diseases

Since the *in-silico* mutagenesis assay implemented in the TransRBP provides an interface for predicting genetic variants’ effect on RBP binding, we next sought to identify disease variants with a molecular mechanism of altering RBP binding. We focused on the genetic variants that are associated with ten brain-, ten heart-, two lung-, and two muscle-relevant diseases collected from ClinVar^62^ (**Fig. 4a, Extended Data Fig. 7a, and Methods**).

**Figure 4.**
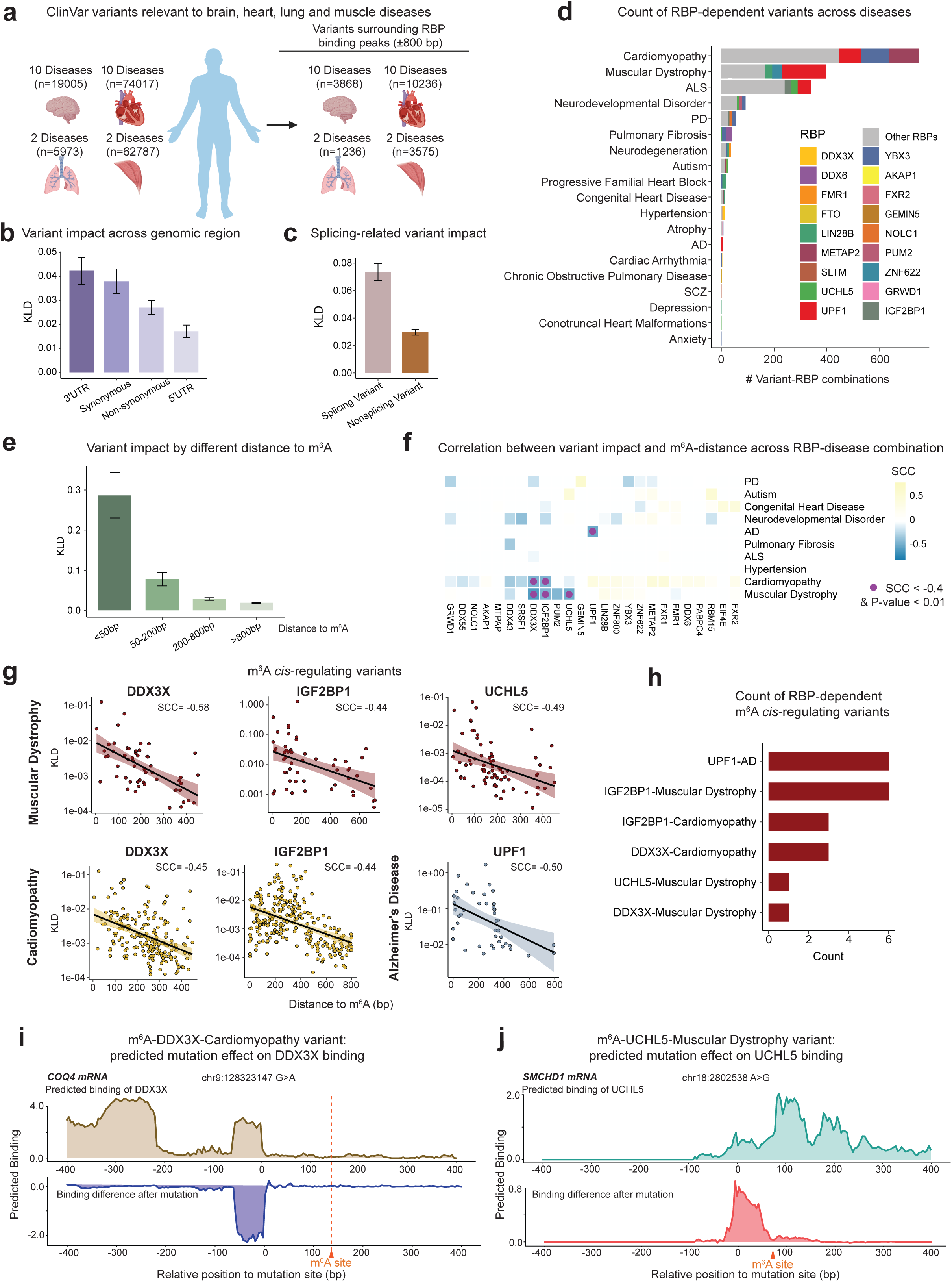
TransRBP interprets disease variants mediated through RBP binding and m^6^A. **(a)** Selection of ClinVar variants from 10 brain disorders, 10 heart disorders, 2 lung disorders, and 2 muscle disorders within 800 bp of the binding sites of RBPs. **(b)** Variant impact (KLD) scoring across 32 m^6^A-related RBPs on different genomic regions. 3’ UTR: 3’ untranslated region. 5’ UTR: 5’ untranslated region. The bars represent the mean values, and the error bars represent the standard errors. **(c)** Variant impact (KLD) scoring across 32 m^6^A-related RBPs for the genetic variants located at the splicing junction and outside the region. The bars represent the mean values, and the error bars represent the standard errors. **(d)** Identification of variants-RBP combinations across genetic diseases where variants strongly alter the RBPs binding. The RBPs with top 3 associated genetic variants in each disease are annotated with colors, and the rest of RBPs are colored in grey. ALS: amyotrophic lateral sclerosis; PD: Parkinson’s disease; AD: Alzheimer’s disease; SCZ: schizophrenia. **(e)** Variant impact (KLD) grouped by the distance of disease variants to their nearest m^6^A sites. The bars represent the mean values, and the error bars represent the standard errors. **(f)** Heatmap showing the Spearman correlation (SCC) between variant impact (KLD) and variant’s distance to m^6^A site across RBP-disease combinations with over 10 variants. Purple dots represent the RBP-disease combinations showing SCC < -0.4 and P-value < 0.01 **(g)** Scatter plot of variant impact (y-axis) against the distance to m^6^A site (x-axis) across 6 significant RBP- disease combinations with SCC < -0.4 and P-value < 0.01. **(h)** Count of RBP-dependent variants within the 6 RBP-disease combinations whose variants impact show m^6^A *cis-*acting effect. **(i)** Example of the predicted mutation effect of a cardiomyopathy variant, chr9:128,323,147:G>A, on the binding of DDX3X to the *COQ4* mRNA. The predicted disruption of RBP binding upon the genetic variant is visualized in the bottom panel. **(j)** Example of the predicted mutation effect of a muscular dystrophy variant, chr18:2802,538:A>G, on the binding of UCHL5 to *SMCHD* mRNA. The predicted increase of RBP binding upon the genetic variant is visualized in the bottom panel.

We assessed the impact of these variants on the binding of each corresponding RBP by KLD via the *in-silico* mutagenesis assay. The genetic variants of muscle-related diseases show globally highest impact on RBP binding, especially for Muscular Dystrophy (**Extended Data Fig. 7b**-c).

Interestingly, the disease variants located in the 3’ UTR region where m^6^A is enriched^46^, exhibited the most pronounced impacts on RBP binding, while the variants in 5’ UTR showed the weakest effects (**Fig. 4b**). Moreover, the synonymous variants possessed stronger effects on RBP binding compared to the non-synonymous ones, indicating a post-transcriptional mechanism for these variants that do not alter protein sequence. Additionally, variants adjacent to the splicing junction showed a globally higher impact on the binding of m^6^A-related RBPs (**Fig. 4c**), in line with our observation that these RBPs are enriched for the binding at splicing consensus. These results collectively provide orthogonal regulatory mechanisms dependent on altering RBP binding and splicing other than protein sequence changes in disease pathogenesis.

To further prioritize RBP-dependent disease variants, we defined 1,806 variant-RBP combinations where the variants are predicted to strongly alter RBP binding (**Fig. 4d**, **Supplementary Table S4**), using the predicted impact of randomly-selected 100 thousand mutations surrounding the binding sites of each RBP as background (**Extended Data Fig. 7d, Methods**). Each of these variants could strongly alter the binding of one or multiple RBPs (**Supplementary Table S4**). For brain disorders, we identified 340 variant-RBP combinations in ALS, 92 in neurodevelopmental disorder, 56 in Parkinson’s disease (PD), 36 in neurodegeneration, 25 in autism, and fewer across Alzheimer’s, SCD, anxiety, and depression (**Extended Data Fig. 7e**). In heart conditions, 750 variants were associated with cardiomyopathy, with smaller numbers in other diseases. For lung disorders, we discovered 40 variants in pulmonary fibrosis and 2 in chronic obstructive pulmonary disease. For muscle disorders, we found 398 variants in muscular dystrophy and 8 in atrophy (**Extended Data Fig. 7e, Supplementary Table S4**). These variants are suggested to lead to the disease pathogenesis via the molecular mechanism of RBP binding disruption in the relevant tissues.

### Identification of disease variants dependent on m^6^A-RBP interaction

m^6^A modification has been revealed to play fundamental roles in various human diseases, such as cancer progression, immune dysfunction, neurological disorders, and more^9–14^. Given our model’s ability in dissecting m^6^A-related RBP binding, we next investigated the mechanistic relationship between m^6^A and genetic disorders mediated through RBPs. Intriguingly, we observed a distance-dependent relationship between m^6^A modification and genetic variants’ impact on RBP binding (**Fig. 4e**), indicating that variants located in the proximity of m^6^A tend to show stronger RBP binding disturbance.

We then searched for the disease variants showing pronounced m^6^A distance-dependent effects on RBP binding (**Methods**). We identified 6 RBP-disease combinations, where the proximity of variants to m^6^A sites shows a strong correlation with the predicted impacts on RBP binding (SCC < -0.4 and P-value < 0.01) (**Supplementary Table S5**). These included the m^6^A-RBP interactions pertaining the binding of DDX3X, IGF2BP1, and UCHL5 with Muscular Dystrophy, DDX3X and IGF2BP1 with Cardiomyopathy, and UPF1 with AD (**Fig. 4f-g**).

Among these m^6^A *cis-*acting regions, we further prioritized 20 disease variants that strongly alter RBP binding based on the *in-silico* mutagenesis (**Fig. 4h, Supplementary Table S5**). For example, we identified a cardiomyopathy-related mutation chr9:128323147:G>A locating at 140 bp to an m^6^A in *COQ4*, which is a gene crucial for coenzyme Q biosynthetic and mitochondrial respiratory chain function^63,64^. We illustrated a potential mechanism that this variant can disrupt the DDX3X binding on *COQ4* mRNA in an m^6^A-dependent manner, thereby contributing to the pathogenesis of cardiomyopathy (**Fig. 4i**). In another example, a muscular dystrophy-associated mutation chr18:2802538:A>G located in *SMCHD1*^65^, was predicted to act as an m^6^A *cis-*acting variant that may strongly impact the binding of deubiquitinase UCHL5 onto *SMCHD1* mRNA (**Fig. 4j**).

Collectively, our results delineate an m^6^A-dependent variant-RBP regulatory axis for various genetic diseases. Taking advantage of our model interpretation assays, we provide a complex disease mechanism that disease variants can alter RBP binding in an m^6^A *cis-*regulating manner.

## Discussion

In this study, we presented TransRBP, an interpretable sequence-to-signal deep learning model that enables *de novo* RBP binding prediction and interpretation from RNA sequences. TransRBP outperformed the state-of-the-art RBP deep learning model and demonstrated accuracy towards experimental replicates. Combining contribution scores, self-attention mechanisms, and in-silico mutagenesis tools, we modeled the binding landscape of 32 m^6^A-related RBPs and disentangled their interaction with m^6^A. Importantly, we shed light on a potential m^6^A-RBP interaction mechanism for genetic disease variants, where variants alter RBP binding in an m^6^A *cis*-acting manner.

The intrinsic similarities between genomic sequences and natural language have paved the way for adapting NLP models including deep learning to genomic studies. While deep learning models have been applied for RBP binding prediction, most were focused on classification-based approaches that determine whether an RBP binds or not^40,41,42^. Our study transcended previous classification-based RBP modeling by predicting RBP binding at base-pair resolution and enabling motif discovery from RNA sequences. Moreover, through multiple innovations in model design and data augmentation strategies, TransRBP outperformed the current sequence-to-signal model by 28% accuracy improvement.

We interpreted the m^6^A-related RBPs binding motif using a gradient-based interpretation method, which is different from traditional tools. Traditional methods primarily identify motifs by detecting statistical overrepresentations against a sequence background^55^. In contrast, our method identified motifs based on their ability to accurately predict experimental binding profiles without explicitly defining sequence features a priori. We noted the enrichment of m^6^A-related RBP motif on splicing consensus, suggesting unexpectedly wider crosstalk between m^6^A and splicing through RBPs.

Moreover, our approach allows for the learning and integration of motif cooperativity and provides a quantitative interpretation of sequence contribution to RBP binding.

Our study demonstrated how *in-silico* modeling can decipher the interaction between two important post-transcriptional regulations – RBP binding and m^6^A modification. An important step towards this was the multiple interpretation tools we established along with the TransRBP model. We innovatively defined the impact score to disentangle the impact between m^6^A and RBP binding with *in-silico* mutagenesis assays and systematically evaluated the effects of mutations near m^6^A sites. Moreover, we leveraged the self-attention mechanism embedded in TransRBP to allow the model to weigh the importance of different positions along the sequence when making predictions, effectively capturing the contextual interactions underlying RBP binding near m^6^A sites. With these approaches, we identified previously unappreciated RBPs that are likely to be influenced by m^6^A.

Our studies provided potential molecular insights into the regulatory underpins of genetic variants at the post-transcriptional level. We assessed the impact of ClinVar disease variants on RBP binding, spanning 24 genetic diseases relevant to brain, heart, lung, and muscle tissues. Importantly, we identified 1,806 variant-RBP combinations, where variants strongly alter the binding of one or multiple RBPs, and thus act as a potential mechanism for genetic disease pathogenesis. Since the genetic regulatory map of RBP binding is currently missing, our approach represents an initial attempt for connecting RBP binding disruption to genetic diseases at the genome-wide scale.

Furthermore, we attributed these variants with dependence on m^6^A-RBP interactions. We found 6 RBP–disease combinations, including DDX3X, IGF2BP1, and UCHL5 with muscular dystrophy; DDX3X and IGF2BP1 with cardiomyopathy; and UPF1 with Alzheimer’s disease. In these cases, the variants’ impact on corresponding RBP binding showed a distance dependency to m^6^A sites, suggesting a *cis-*acting role of m^6^A on disease variant impact associated with RBP binding.

Despite the prominent performance of TransRBP and its unique biological and pathological insights, our study has several limitations. First, while Transformer may implicitly extract information on RNA structure for prediction, TransRBP does not explicitly account for RNA secondary and tertiary structures, which are known to influence RBP binding and may interact with m^6^A modifications. We envision that incorporating structural information can further enhance the model’s predictive accuracy and potentially identify crosstalk between structures, RNA modifications, and RNA sequences on RBP binding. Second, despite the data augmentation strategy we developed for increasing the prediction power, the model’s accuracy and interpretability can be further promoted with a substantially increased quantity and diversity of the training data.

In conclusion, TransRBP not only represents a robust framework for predicting RBP binding and enabling reliable motif discovery, but also acts as an interface for RBP-related biological interaction and genetic disease mechanism interpretation. We anticipate that our model will facilitate our understanding of functional genomics and shed light on post-transcriptional regulatory mechanisms in the field of disease genetics.

## Methods

### eCILP data collection and processing

The enhanced CLIP (eCLIP) data from the K562 cell line used as RBP binding profiles in this study was obtained from publicly available sources at ENCODE^44,45^ (www.encodeproject.org/). In total, we curated 139 RBP datasets, each of which contains two replicates of an eCLIP experiment and a size-matched input (SMInput) experiment serving as a critical control for nonspecific background signals (**Supplementary Table 1**). The experiment data of 115 out of 139 RBPs were paired-end (PE) sequenced and the rest were single-end (SE) sequenced, both aligned to the hg38 human reference genome. For those PE eCLIP and SMInput data, only the second read in each pair was kept for use via samtools^67^ (v.1.5) to avoid potential bias caused by sequencing strategies. We next merged two eCLIP replicates and paired them with the corresponding SMInput using log2 fold- change with a bin size of 1 and excluded the K562 blacklist regions via deepTools^68^ (v.3.5.4) to mitigate experimental biases. The single-base RBP binding sequencing coverage values extracted from the processed BigWig files were used as the ground truth of TransRBP training.

### Preprocessing and peak calling of MeRIP-seq data

MeRIP-seq data of K562 cells was acquired from Gene Expression Omnibus with accession number GSE137752. Raw sequencing data was aligned to the hg38 reference genome by STAR^69^ (v.2.7.11) and the peaks were called by macs2^70^ (v.2.2.6) using the matched RNA-seq data as control. The narrowPeaks called from macs2 was used for further analysis.

### RNA sequences for TransRBP

We used the hg38 human reference genome sequence from the UCSC Genome Browser database^71^. For each RBP, we extracted their binding peak information, including chromosome, start site, end site, and strand from narrowPeak files downloaded from ENCODE, which have been normalized and analyzed for irreproducible discovery rate (IDR), considering both eCLIP replicates and their SMInput. Here we utilized a method of data augmentation to acquire RNA sequences for TransRBP (**Extended Data** Figure 2c) and avoid the potential model fitting to conservative RBP binding shape and increase the data diversity. This is achieved by using an 800-bp sliding window that moves throughout each peak, with a step size of one-tenth of the RBP peak length. The start sliding extended half of the window length (400bp) outwards from the start site of the peak. The RNA regions that overlap with peaks were treated as positive samples and the rest as negative samples. In this way, the ratio of positive vs negative samples was approximately kept to 2:1, and more RNA sequences were obtained around the 5’ end of peaks, as they have been previously reported more likely to hold crosslinking sites^43,66^. To further mitigate the risk of overfitting to particular RBP binding profiles and to control for performance biases due to variable data volumes across RBPs, we randomly selected 20,000 sequences (both positive and negative) for each RBP to form the dataset.

### TransRBP architecture

TransRBP takes obtained one-hot-encoded RNA sequence (A=[1,0,0,0], U=[0,1,0,0], C=[0,0,1,0], G=[0,0,0,1], N=[0,0,0,0], respectively) as input to predict RBP binding profiles at single-base resolution as output. TransRBP employs a convolution-attention architecture that consists of an encoder, a transformer module, and a decoder (**Extended Data** Figure 1d). The architecture was implemented in Pytorch^72^ (v.2.0.1).

TransRBP starts with a 1D convolution layer with 256 filters of kernel size = 5, padding = 2, and stride = 1, a batch normalization layer, and an exponential activation, to transform the input dimension from (N × 4 × 800) to (N × 256 × 800), where N is the batch size. The encoder contains two residual blocks, each of which is composed of (1) a 1D convolution layer with 256 filters of kernel size = 3, (2) a batch normalization layer, and (3) a ReLU activation. The encoder helps promote information propagation and robust motif derivation from gradient-based feature attribution.

The transformer module is built with 4 transformer encoder layers. We set the number of hidden layers to 256, the number of attention heads to 8, and the dropout rate to 0.1. We applied the sinusoidal key-query positional embedding and set the maximum length to 800. The transformer module concatenates each position of the input sequence to every other position to form an 800-by- 800 interaction map via the self-attention mechanism. The transformer module helps learn long-range and high-order interactions among sequences.

The decoder contains four residual blocks, each of which is composed of (1) a 1D dilated convolution layer with 256 filters of kernel size = 3, (2) a batch normalization layer, and (3) a ReLU activation. We designed the dilation at the corresponding layer to be 2, 4, 8, and 16. The decoder helps increase the receptive field and reinforce organizations between different elements.

TransRBP ends with a 1D convolution layer of kernel size = 1 to combine 256 channels down to one channel and transform the features to output (N × 1 × 800) as the predicted RBP binding profile.

### TransRBP training and hyperparameter tuning

RBP binding profiles from eCLIP experiments in K562 cells paired with the corresponding SMInput experiments were used to train TransRBP. For each RBP, a total of 20,000 800-bp RNA sequences were randomly selected after data augmentation, and were then split by chromosome into validation (chr2, chr3, chr4), test (chr1, chr8, chr9), and training (all other autosomes) sets in a ratio of 2:1:7.

The training of TransRBP used a batch size of 64, the Adam optimizer with an initial learning rate = 1e-3, the cosine annealing learning rate scheduler, the gradients clipping by max norm below 5, the maximum epochs of 100, and early stopping with the patience of 10 epochs. The mean squared error (MSE) loss was calculated to compare the predicted and the experimental profiles (ground truth). The Pearson correlation coefficient (PCC) and the Spearman correlation coefficient (SCC) were calculated as evaluation metrics. To ensure the performance of the model, we randomly selected 8 out of 32 m^6^A-related RBPs and carried out a comprehensive hyperparameter assessment, together involving the learning rate, the number of encoder layers, and the number of transformer layers. The hyperparameters were selected based on an overall assessment of PCC, SCC, and MSE on their test sets (**Extended Data** Figure 2a-b).

### TransRBP evaluation and benchmarking

To evaluate predictive performance, we systematically compared the predicted RBP binding profile versus two negative controls across all RNA sequences in the test set based on PCC and SCC: (1) the SMInput background signal for each eCLIP experiment, and (2) the profile predicted by naive TransRBP model with the same architecture but randomly initialized parameters. The correlation between two eCLIP experimental replicates (normalized by SMInput with log2 fold-change) on the test set for each RBP was also calculated.

We then carried out benchmarking by comparing TransRBP with the state-of-the-art sequence-to-signal model, RBPNet^43^. We utilized the same data augmentation method and selected the same number of training samples as TransRBP. RBPNet was implemented using its training strategy as mentioned in the original paper, including the hyperparameter settings, the negative log-likelihood as the loss function. We also used PCC and SCC separately as evaluation metrics on test sets to compare the performance.

### Integrated gradient contribution scores calculation

Integrated gradient (IG)^52^ is a deep learning interpretation algorithm that attributes the prediction of a deep learning model to the importance of input features. RNA sequences with high contribution scores positions are recognized as bases important for RBP binding signal prediction. IG algorithm calculates and integrates the gradients of the model’s output along the path from the neutral baseline (here we use zero-vectors) to the input:

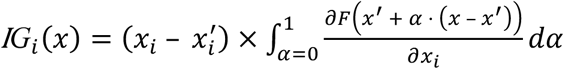

where ***F*** is a scalar neural network function, ***x*** is the actual input to the model, ***x′*** is the baseline input, and ***α*** scales the difference between the baseline and the input, ranging from 0 to 1.

By default, IG attributes the contribution score of a deep neural network to a model prediction of scalar output. This naturally holds for classification-based tasks where the output is usually the probability for classification. However, the output of TransRBP is not a scalar, but a 1D tensor containing the predicted binding signal on each base. We, therefore, adapted the idea in multiclass classification and transformed the output into a scalar by a weighted sum of probabilities, which is in analogy to Avsec *et al*.’s generalization of DeepLIFT^36^ :

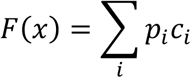

where *c*_i_ is the predicted signal of each position and ***p**_i_* is the probability values acquired by normalizing the profile predictions using the softmax function ***p**_i_*=softmax(x)*_i_*. This ensures that positions with high predicted values are given more weight and thus recognized as more important. To avoid extensive backpropagation resulting from the positional probability weighting, we mutate the function to treat the probability value ***p**_i_* as a constant:

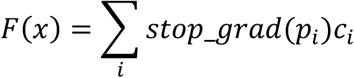

where the *stop_grad* function represents the probability value disabled from gradient calculation. The IG function is implemented based on captum^73^ (v.0.6.0).

### Gradient correction

To further improve the interpretability of our gradient-based IG algorithm, we applied a simple statistical gradient correction method presented by Majdandzic *et al.*^53^, to account for removing spurious noise in the gradients. Given the input of RNA is a one-hot encoded simplex vector that accounts for 4-categories categorical data, gradient correction removes the off-simplex gradient component orthogonal to the direction of the simplex:

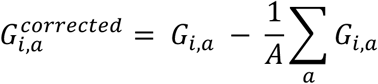

where *i* is the position in the input, *a* is the index representation of 4 types of nucleotides, and *A* is the number of categories (here *A*=4, standing for 4 classes of nucleotides). In short, the gradient correction subtracts the mean gradient of position *i* from the original gradients of each of the 4 nucleotides on the corresponding position.

### Motif discovery and clustering from IG scores

We applied a 5-mer scanning window on 800 nt surrounding the 5’ end of each ENCODE narrowPeak, the position reported to more likely harbor the binding peak^43,66^, then summing the contribution scores of each 5-mer. To prioritize high contribution motifs and account for the differential peak number of each RBP, the top 10,000 highest sum of contribution scores of 5-mers were selected for each RBP and used for clustering.

To cluster motifs based on the similarity and account for potential truncation events in the motif, we allowed the 5-mers to be in the same cluster if they only harbor one mismatch or could be matched with one shift. For each RBP, we started off the clustering from the 5-mer who has the highest contribution scores and identified all other 5-mers within the range of one mismatch or one shift.

Once these identified 5-mers were considered in one cluster, they will be removed and not considered for further clustering action. The process then iterates to start clustering with the highest contribution 5-mer remaining in the 10,000 5-mers list and ends until all the 5-mers are clustered.

The clustered 5-mers were then placed on a 7-mer position considering those who matched with one shift. The position weight matrix (PWM) was derived for each cluster based on nucleotide frequency and used as a representation for the motif. These motifs were ranked based on the number of supporting 5-mers involved in constructing the cluster. In other words, clusters with more supporting high-contribution 5-mers were prioritized as a better representation of the motifs of RBP.

The top 3 PWMs gathered from the clusters were then selected as the motifs for each RBP and were further hierarchical clustered based on their similarity using motifStack^74^ (v.1.42.0) with cutoffPval = 0.01.

### Motif network analysis

The RBP motif network is constructed based on the motif matching from the identified top 3 motifs across the 32 m^6^A-related RBPs and motif reference from ATtRACT^56^ database. Given the variability in motif lengths within the ATtRACT database, our matching criteria included exact matches to the query 5-mers and instances where the query 5-mers were contained within longer motifs. This approach ensured a thorough and relevant motif comparison. In the network presented in Figure 2e, the width of the edge represents the number of matched motifs between RBPs, while the node size is the sum of all matching events related to an RBP. Larger nodes signify a higher degree of motif similarity with the set of 32 m^6^A-related RBPs, suggesting key players in the motif- based interaction of these RBPs.

### *In-silico* mutation impact scoring

TransRBP predicts RBP profile solely based on RNA sequence, thus disruption of crucial sequence can have a significant impact on RBP binding. For each of the variants derived from *in-silico* mutagenesis, we estimate its impact by the change of RBP binding profile distribution of the alternative allele (alt) compared to the reference allele (ref)^43^. We defined the change of distribution of a given point mutation by the KL divergence of the alternative allele compared to the reference allele, given by:

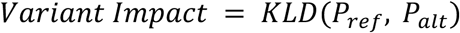

where *P_ref_* and *P_alt_* are the binding probability distribution normalized from the predicted profile of the reference allele sequence and alternative allele sequence

### Sequence-based RBP-m^6^A interaction analysis

MeRIP data peaks were acquired from preprocessing (see above) and were intersected with the binding peaks of each RBP. For these intersected MeRIP peaks, the submit (maximum) locations of the m^6^A footprint were then determined as the center for 400-bps flanking *in-silico* mutagenesis. We performed the saturated in-silico mutation on the 800 bps sequence surrounding the putative m^6^A site, where each of the original nucleotides is mutated to the other three alternative nucleotides.

We defined the m^6^A to RBP impact by the differential impact of mutation proximal to the m^6^A site (<50 bps) and the mutation distal to the m^6^A site (>50 bps). To this end, distal mutation impact serves as a background that varies with each RBP model and thus enables comparison across RBPs. The m^6^A-to-RBP impact was quantified using the ratio of mean variant impacts between proximal and distal mutations:

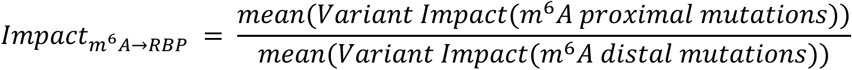

### Attention-based RBP-m^6^A interaction analysis

The attention map within the Transformer illuminates the intensity of interactions between positions within the sequence^48^. The attention weights at position (i, j) indicate how much the i-th element in the output (RBP profile) should consider the j-th element in the input (RNA sequence). To further analyze m^6^A-to-RBP impact, we exploited the self-attention mechanism embedded in the Transformer to model the interaction.

We acquired the MeRIP peak submit site that intersected with each RBP peak and extended 400 bp flanking region as inputs to the model, and acquired the mean attention weight across the inputs from Transformer. For each input, the attention weight is gathered by collapsing the attention heads to a single L by L matrix by taking the maximum at each position as suggested by Ullah *et al.*^48^.

This method aggregates the self-attention outputs from the multiple subspaces, each associated with individual heads, into a unified attention profile. This condensed view helps in summarizing and analyzing the dominant attention patterns more effectively. Then the attention weights of these inputs are subtracted by the mean attention weights of 2,048 random input sequences to serve as a way of normalization.

Since there is no max-pooling in the encoder of TransRBP, the sequence position in input is preserved to the self-attention. Thus, the attention weight indicates the importance of the RNA sequence in input when making output RBP profile predictions. We placed all the putative m^6^A modifications at the center of the input sequence. High attention scores at this position suggests a substantial impact on the output RBP profile in the position of m^6^A, thereby indicating a m^6^A-to-RBP interaction.

### Identification of RBP-dependent disease variants

We obtained genetic variants associated with specific diseases from ClinVar, focusing on those relevant to 10 brain diseases, 10 heart diseases, 2 lung diseases, and 2 muscle diseases. To specifically focus on RBP-related variants, we selected variants located within 800 base pairs of binding peaks of 32 m^6^A-modified RBPs (**Supplementary Table S4**).

We then evaluated the impact of these disease variants on the binding of RBP through in-silico mutagenesis. The impact of each mutation was quantified by KLD (see above). To establish a benchmark for substantial disruption of RBP binding, we generated a background mutation effect by profiling the impact of 100,000 randomly selected point mutations, within 800 base pairs of RBP binding peaks. We then select the mutation impact of the top 10% for each RBP as a cutoff to define the variant’s function potential through a mechanism of distorting RBP binding.

### Discovering m^6^A *cis*-acting effect on variants

To investigate the role of m^6^A modifications in mediating the effect of disease-associated variants on RBPs, we calculated the distance between each variant and its nearest m^6^A site. We only focus on variants within 800 bps of m^6^A to ask for a *cis-*regulatory role of m^6^A. Variants located within 800 base pairs of an m^6^A site were considered for analysis to define a potential *cis-*regulatory role of m^6^A in modulating variant effects. To maintain statistical power and avoid strong correlations arising from sparse data, we limited our analysis to RBP-disease combinations containing at least 10 variants.

We define the *cis-*acting function of m^6^A as a distance-dependent role in influencing the impact of disease variants on RBP binding. In other words, variants more proximal to m^6^A have a stronger effect in disrupting RBP binding. We then quantified the correlation between the distance of each variant from the nearest m^6^A site and its impact on RBP binding for each RBP-disease combination. This correlation was measured using SCC (or Spearman’s R). RBP-disease combinations with a SCC < -0.4 and P-value < 0.01 were selected, identifying 6 such combinations. m^6^A modifications in these combinations are suggested to contribute to the variant-induced disruption of RBP binding in a distance-dependent manner.

## Acknowledgements

We thank the members in the Xiong Lab for discussion and suggestions throughout the project. We thank the support from the core facilities and computing platform of Liangzhu Laboratory at Zhejiang University. This work was supported by the National Natural Science Foundation of China (nos. 32422017, 32370609 and 92353301 to X.X.), Ministry of Science and Technology of China (nos. 2024YFF1207600 and 2023YFA1800700 to X.X.), and the funding from Liangzhu Laboratory at Zhejiang University and the State Key Laboratory of Transvascular Implantation Devices to X.X..

## Author contributions

This study was designed by J.L., X.Z., Y.Y. and X.X., and directed and coordinated by X.X.. X.Z. and J.L. trained and fine-tuned the model with help from Y.Y. Z.X. and J.H., and under the supervision of X.X.. J.L. performed model interpretation analysis under the supervision of X.X.. All authors participated in the discussion of the project. J.L., X.Z. and X.X. wrote the manuscript.

## Code availability

Code related to the work will be publicly available upon publication.

---

**Extended Data Figure 1 related to Figure 1.**
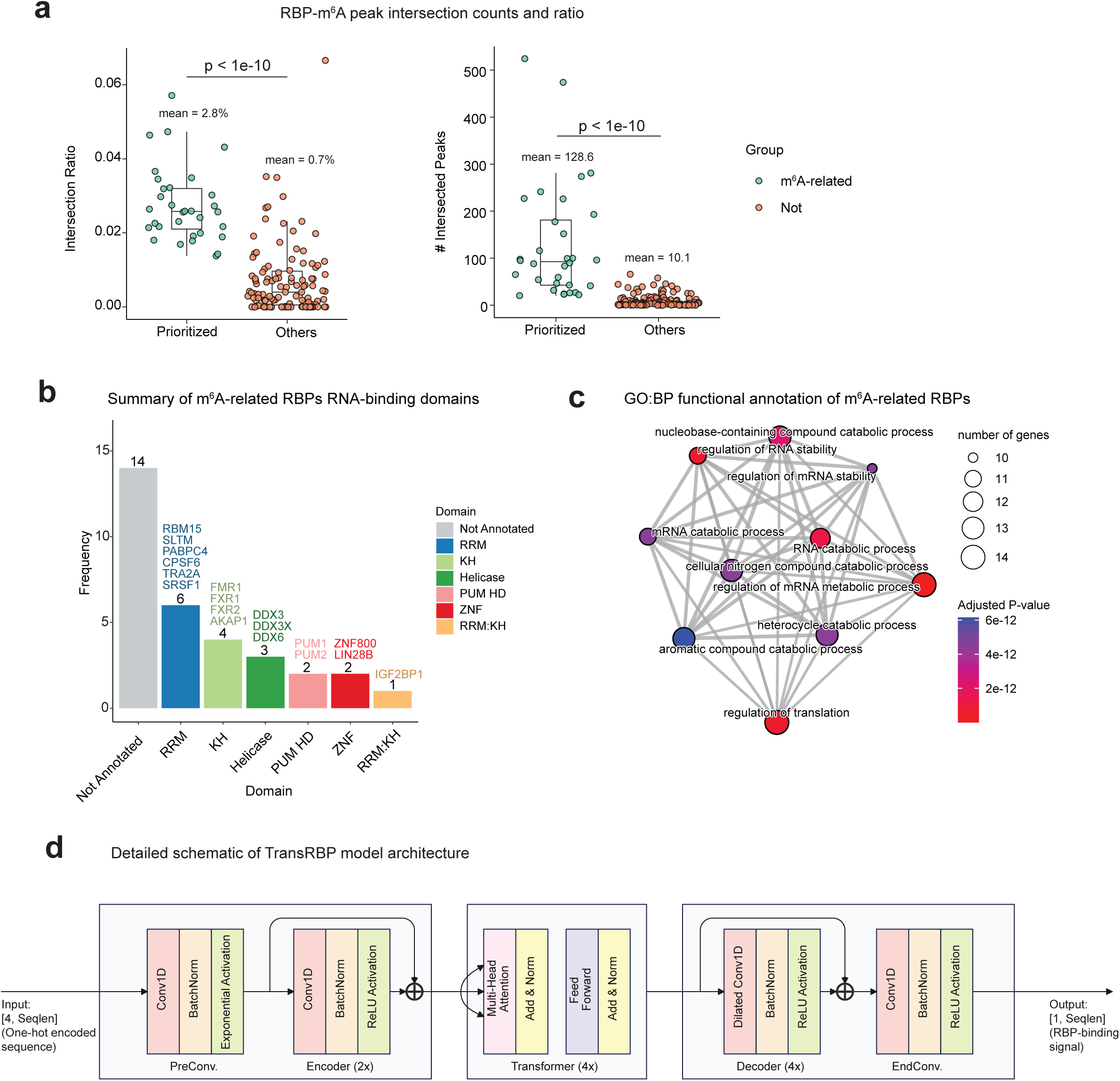
Annotation of m^6^A-related RBPs and TransRBP model architecture. **(a)** Number and proportion of m^6^A-RBP peak intersections between prioritized m^6^A-related RBPs and non-related RBPs. P-values were calculated from the one-way Mann-Whitney U test. **(b)** 32 m^6^A-related RNA-binding domain annotation derived from Nostrand *et al.*^45^. **(c)** 32 m^6^A-related RBP functional annotation from Gene Ontology: biological process (GO:BP). **(d)** Detailed schematic summary of TransRBP model architecture.

**Extended Data Figure 2 related to Figure 1.**
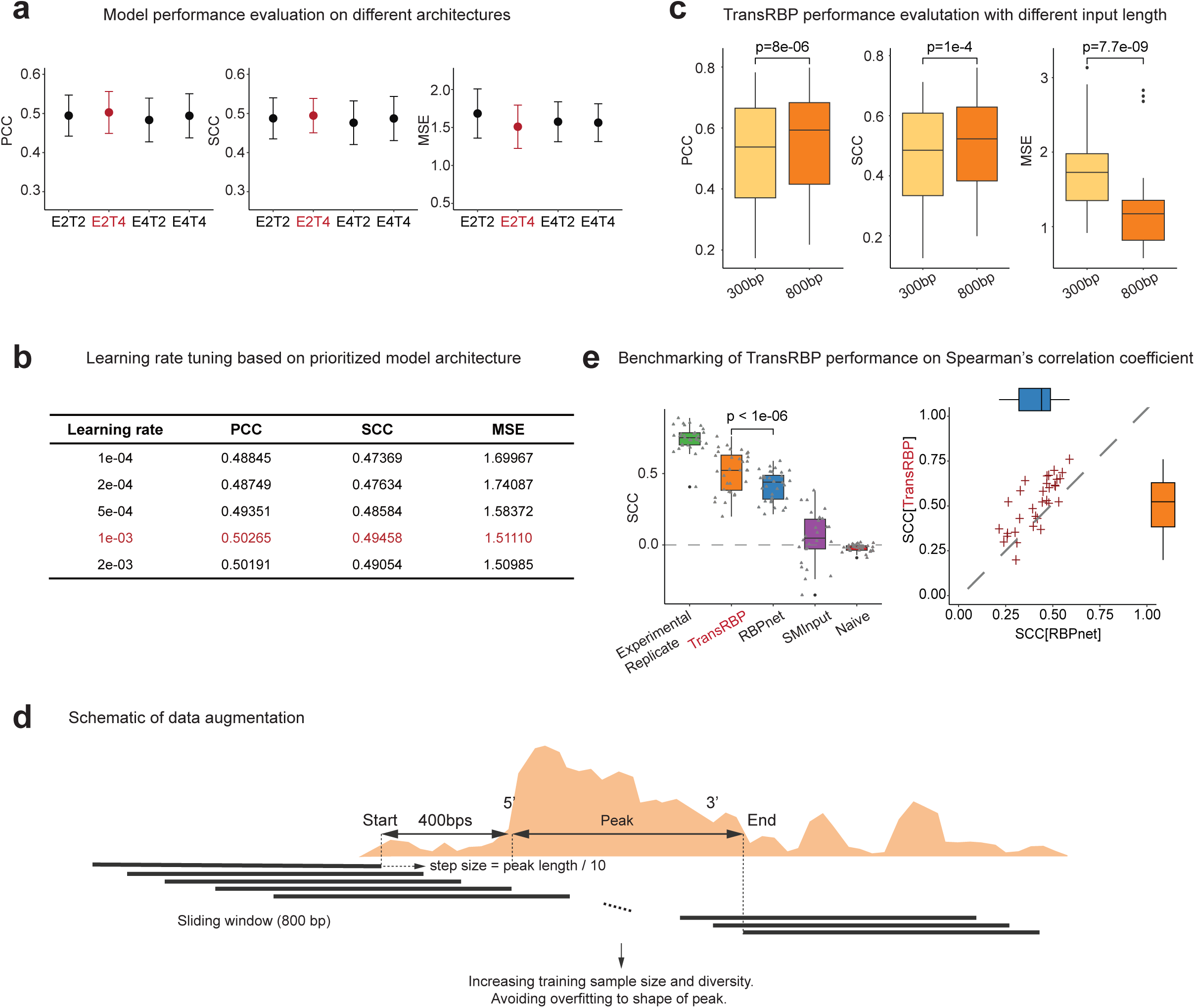
TransRBP model tuning, performance evaluation and strategy of data augmentation. **(a)** Model performance evaluation on different architectures by Pearson correlation (PCC) (left) Spearman correlation (SCC) (middle), and mean squared error (right) on 8 randomly sampled RBPs. E: Encoder layer. T: Transformer layer. **(b)** Table showing the performance evaluation of different learning rates tested on prioritized (2 Encoder + 4 Transformer) architecture on 8 randomly sampled RBPs. **(c)** TransRBP performance evaluation of different input lengths by Pearson correlation (PCC) (left) Spearman correlation (SCC) (middle), and mean squared error (right) on 8 randomly sampled RBPs. **(d)** Schema of data augmentation for inputs. An 800 bp window is placed at a 400 bp 5’ upper to the binding peak, and sliding towards the end of the peak with the step size varying to the peak length. After sliding, a total of 20,000 samples are randomly selected for each RBP. This strategy augmented the RBP peak data as follows. 1. The input sample size is kept the same across RBPs. 2. The ratio of positive vs negative samples is approximately kept to 2:1. 3. More samples are derived from the 5’ end of peaks, previously identified to harbor the binding site^43,66^. 4. the diversity of samples is increased, thus avoiding model overfitting to the shape of peaks. **(e)** Spearman correlation (SCC) between the observed and predicted RBP binding profile evaluated on the held-out test set for TransRBP and RBPnet^43^, and compared to experimental replicates, size-matched input (SMInput) and randomly initialized untrained model (Naive).

**Extended Data Figure 3 related to Figure 2.**
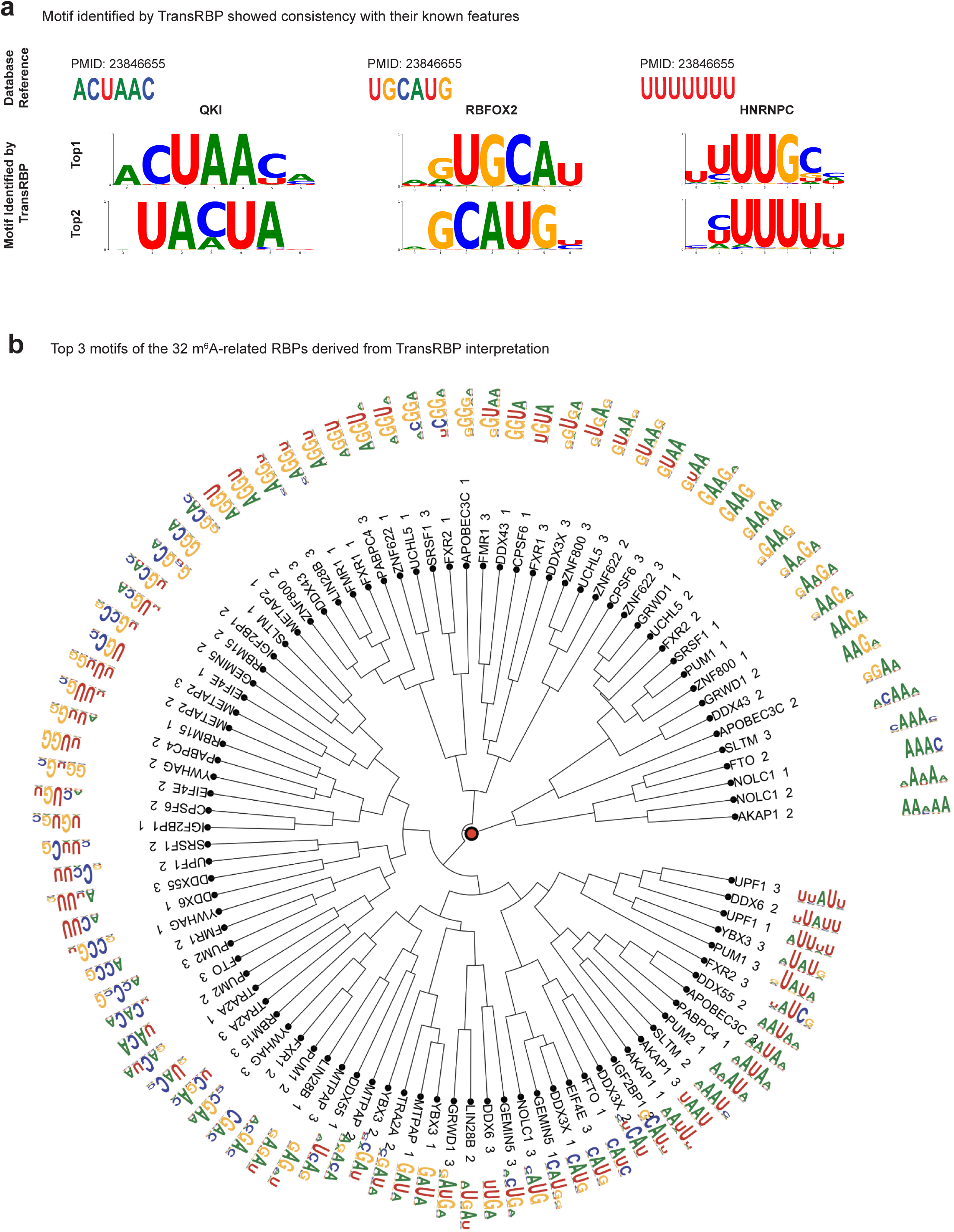
Validation of TransRBP interpretation for motif discovery. **(a)** Gradient-based interpretation of TransRBP recapitulated established motif instances of QKI, RBFOX2, and HNRNPC. Established motif references are shown at the top of each panel. **(b)** Hierarchical clustering plot showing all the top 3 motifs of 32 m^6^A-related RBPs.

**Extended Data Figure 4 related to Figure 2.**
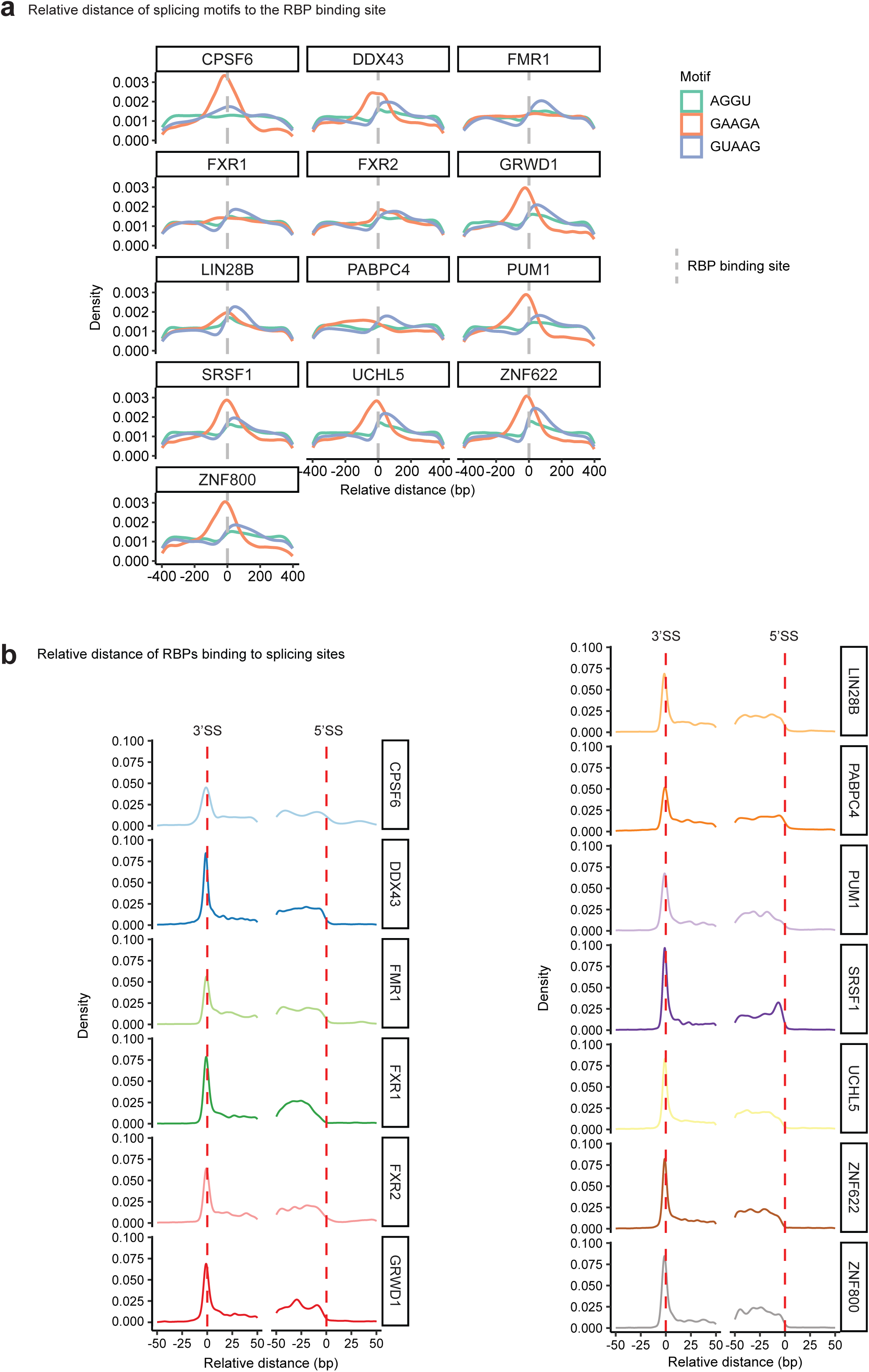
Splicing-related motifs and RBP binding distribution to splicing sites. **(a)** Relative distances of *GAAGA*, *AGGU*, and *GUAAG* sequence motifs to the binding sites of each of 13 splicing-related RBPs. **(b)** Relative distances of the binding sites of each of the 13 splicing-related RBP to splicing sites (SS) on the forward strand.

**Extended Data Figure 5 related to Figure 2.**
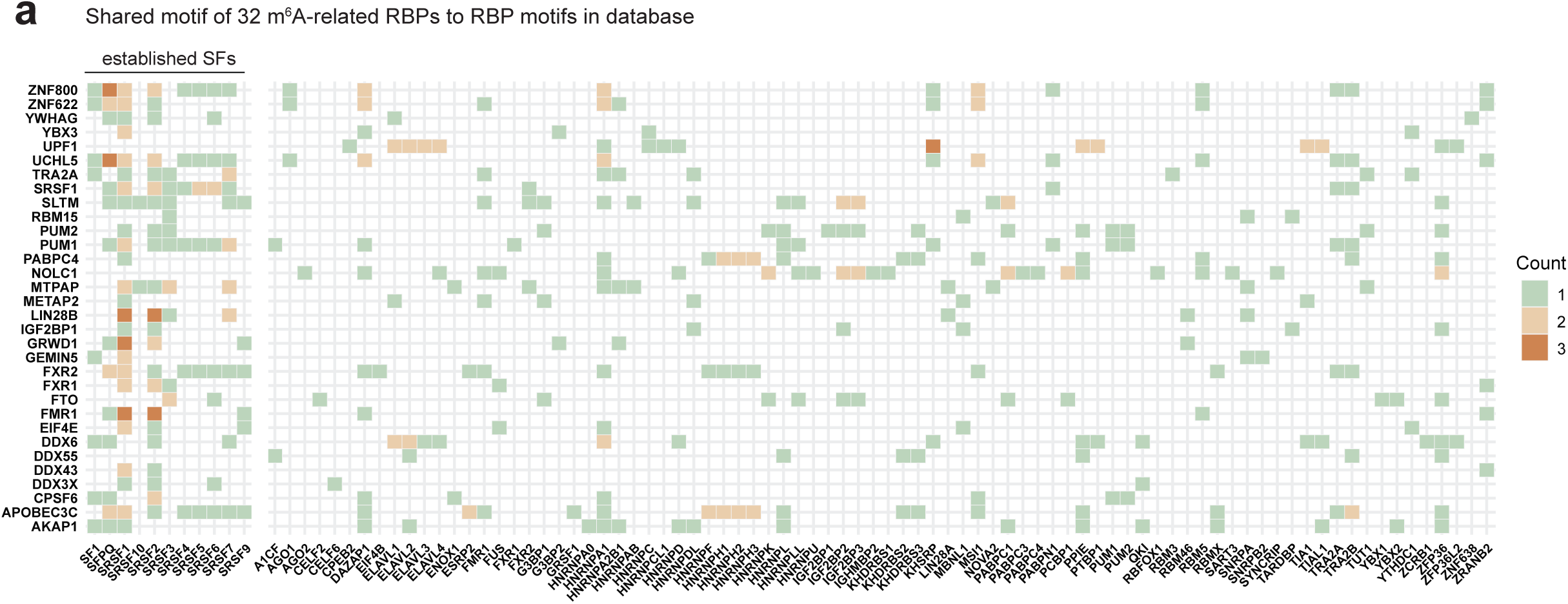
Motif similarity heatmap. **(a)** Heatmap showing the number of shared motifs between top 3 motifs of 32 m^6^A-related RBPs discovered by TransRBP (y-axis), and RBP motifs in the database^56^ (x-axis). Color of the heatmap indicates the number of the shared motifs for each pair of RBPs. SF: splicing factor.

**Extended Data Figure 6 related to Figure 3.**
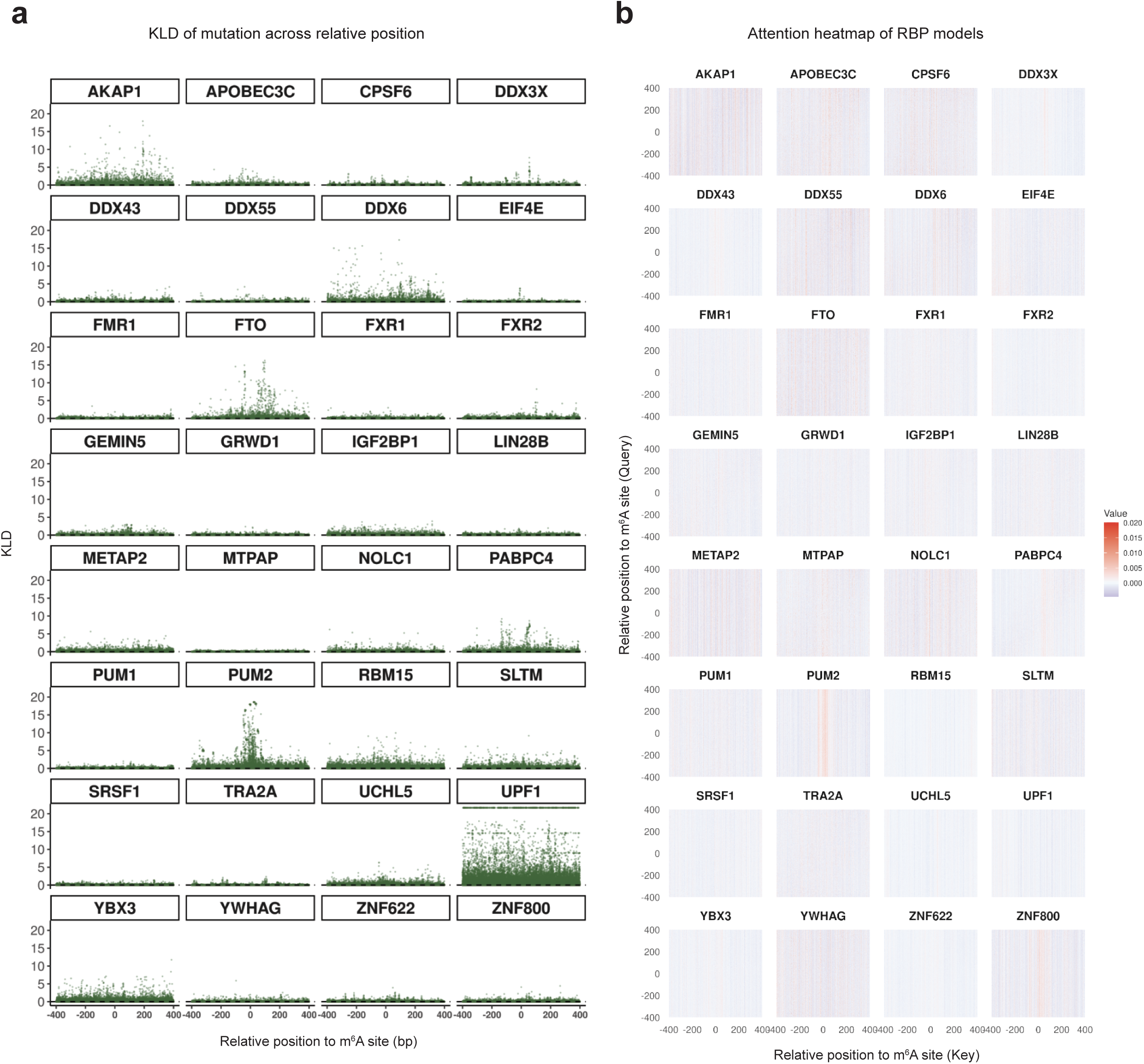
*In-silico* mutation analysis and attention heatmap across 32 m^6^A-related RBPs. **(a)** Distribution of mutation impact (KLD) on RBP binding before and after saturated *in-silico* mutagenesis surrounding m^6^A sites (800bp) for each of the 32 m^6^A-related RBPs. **(b)** Transformer-derived attention heatmap for sequence-to-signal prediction surrounding m^6^A sites (800bp) for each of the 32 m^6^A-related RBPs.

**Extended Data Figure 7 related to Figure 4.**
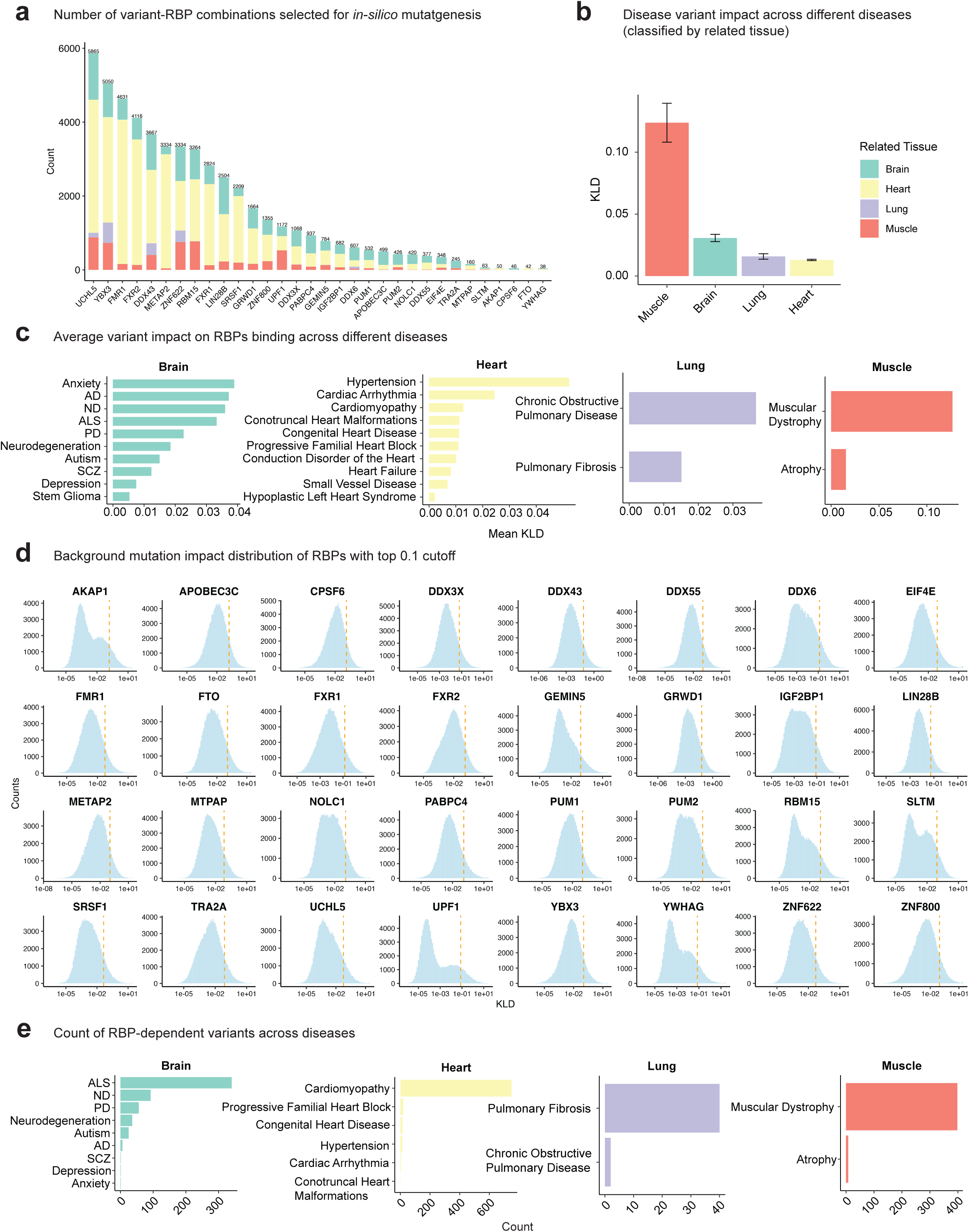
TransRBP enabled disease variant interpretation. **(a)** Count of ClinVar variants within 800 bps of binding peaks for each of RBPs, defined as variant-RBP combinations. Colors indicate the different tissues that diseases are related to. **(b)** Summary of variants’ impact on RBP binding across diseases related to different tissues. The bars represent the mean values, and the error bars represent the standard errors. **(c)** Summary of average disease variant impact on RBP binding across different genetic diseases. **(d)** Distribution of the *in-silico* mutagenesis impact on RBP binding of 100,000 randomly selected single-nucleotide mutations within 800 bps of each of RBP binding sites. The threshold was selected by the top 10% KLD cutoff for each RBP to define the candidate variants that strongly influence the corresponding RBP binding. **(e)** Count of variants that strongly alter the RBPs binding across genetic diseases. ALS: amyotrophic lateral sclerosis; PD: Parkinson’s disease; ND: neurodevelopmental disorder; AD: Alzheimer’s disease; SCZ: schizophrenia.

